# Interaction Between *Yersinia pestis* Ail Outer Membrane Protein and the C-Terminal Domain of Human Vitronectin

**DOI:** 10.1101/2024.01.07.574511

**Authors:** Laurine Vasseur, Florent Barbault, Antonio Monari

**Affiliations:** Université Paris Cité and CNRS, ITODYS, F-75006 Paris, France

**Keywords:** Molecular Dynamics, Bacterial Membrane Protein, *Yersinia Pestis*, Vitronectin, Immune System Evasion

## Abstract

*Yersinia pestis*, the causative agent of plague, is capable to evade human immune system response by recruiting the plasma circulating vitronectin proteins, which acts as a shield and avoids its lysis. Vitronectin recruitment is mediated by its interaction with the bacterial transmembrane protein Ail, protruding from *Y. pestis* outer membrane. By using all atom long-scale molecular dynamic simulations of Ail embedded in a realistic model of the bacterial membrane, we have shown that vitronectin forms a stable complex, mediated by interactions between the disordered moieties of the two proteins. The main amino acids driving the complexation have also been evidenced, thus favoring the possible rational design of specific peptides which, by inhibiting vitronectin recruitment, could act as original antibacterial agents.

**TOC ABSTRACT:** 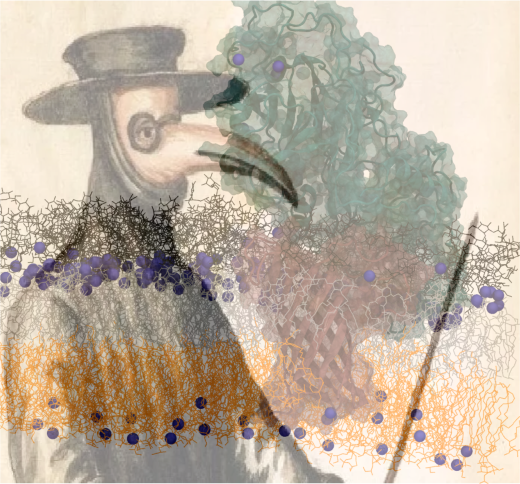

---

Plague is a bacterial infection characterized by a very high mortality rate,^1,2^ which has known recurrent outbreaks significantly impacting public health and the social organization of different countries.^3,4^ As an example, one of the first recorded pandemics, the so-called Justinianic plague has striken the byzantine empire under Justinian rule and between 541-750 a.c., and has also caused significant social, economic, and cultural changes in late antiquity.^5^ The epidemic outbreak starting in 1347, and usually referred as the black death,^6^ is supposed to have caused the loss of one third of the total European population. Other important pandemic events have been recorded during the human history, such as in the 18^th^ century Europe, striking notably England and Northern Italy.^7^ However, less dramatic events have been registered recurrently up to the 20^th^ and the 21^st^ centuries.^8,9^ Furthermore, to date very few treatments are available against the plague,^10^ especially in advanced phases, thus leaving the societies still highly vulnerable against its possible occurrence. Indeed, plague has been classified among the principal emerging infectious diseases, susceptible to cause significant public health and societal threats. Presently up to 600 plague cases are recorded yearly, including occurrences in mainland China, United States, and Brazil testifying of its potential global threat.

The causative agent of plague has been identified in the XIX^th^ century by Alexandre Yersin as a Gramm-negative bacteria, subsequently named *Yersinia pestis*.^11–14^ Rodents and small mammals are efficient secondary vectors of *Y. pestis*, which is primarily spread by fleas’ bites.^15^ However, human-to-human infections due to the inhalation of contaminated air, issued from coughing or breathing, are also common. Different forms of plague, also characteristic of different infection time-scales have been identified. This includes the so-called bubonic plague, mainly affecting the lymph nodes, which become purulent and swollen, thus leading to the appearance of the characteristic buboes; the pneumonic stage^16^ affecting the lungs and significantly reducing the breathing capacity; and, finally, the septicemic plague,^1^ whose outcome is usually fatal, and to which the term of black plague has been associated. This is mainly due to the global darkening of the patient tissues, which happens prior to their necrosis. Furthermore, some instance of antibiotic resistance have also been reported for some *Y. pestis* strains.^17^

*Y. pestis* microbiology has been intensely studied in the past, also at a biophysical and molecular level.^14,18,19^ Recently, the role of the outer trans-membrane Ail protein has been particularly pinpointed.^20–25^ Indeed, Ail being exposed to the outer surface of the bacteria confers *Y. pestis* aggregation capacity,^26^ and participates to the formation of bacterial biofilms and colony,^27^ including in the confined space of human lungs. The structure of Ail in different membrane environments has also been resolved.^23,28,29^ Furthermore, it has been recently evidenced that Ail is also favoring the evasion of the immune system response against *Y. pestis*, thus acting as a strong virulence factor. In particular it has been shown that Ail is able to recruit the plasma circulating human Vitronectin,^30–32^ by interacting with its C-terminal domain.^33,34^ In turn, the bacterial-bound Vitronectin acts as a shield which prevents the complemented-mediated lysis of the bacteria, thus increasing its infectious capacity and virulence.^35^

While a consistent corpus of biochemical evidences indicating the binding of Vitronectin to Ail have been presented,^33,34^ the protein-protein complex has not been characterized at a molecular resolution, and thus, on the one side, the molecular factors favoring the binding have not been elucidated, and, on the other side, no possible rational drug-design strategy aimed at disrupting the Ail/Vitronectin complex, and hence reinstate the immune response, can be envisaged.

The transmembrane part of Ail (Figure 1) is constituted by a rigid β-barrel unit which is flanked, on the outer membrane side, by flexible arms and loops protruding in the polar head and sugar region of the outer membrane lipopolysaccharides.^29^ Interestingly, while the structure of Ail expressed in model lipid nanodisks has been resolved,^29^ and the rational engineering of hyperstable proteins has been proposed,^36^ its behavior on the complex environment of the bacterial outer membrane has not been characterized yet. Conversely, The C-terminal domain of Vitronectin presents a rather rigid core, which is flanked by disordered and non-structured regions.^33,37–39^ As it is common in protein/protein interactions the presence of disordered regions may act as a hub to favor the complexation due to the greater flexibility and structural adaptability.^40–43^

**Figure 1.**
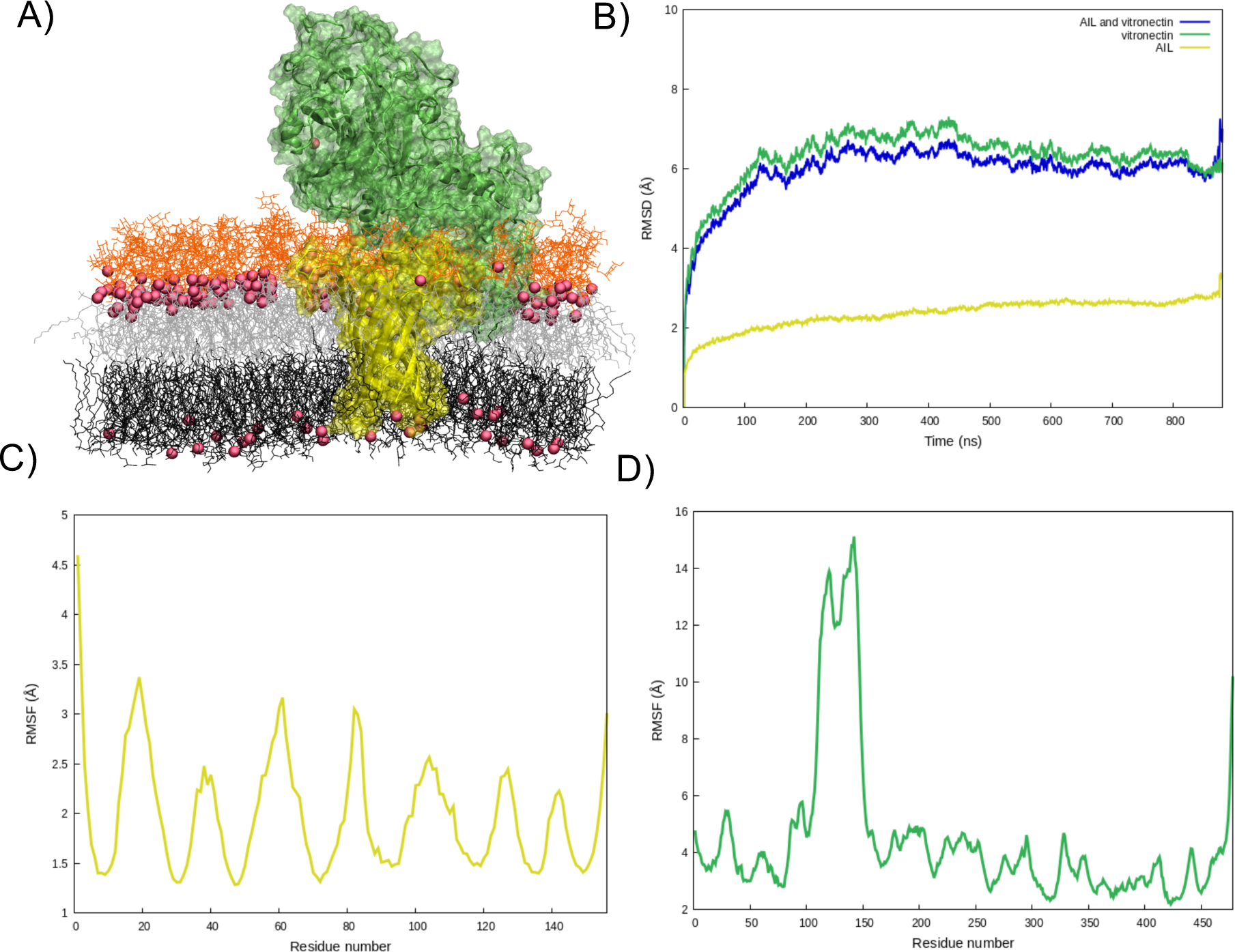
A) Representative snapshot of the Ail/Vitronectin complex embedded in the *Y. pestis* outer membrane environment. The bacterial protein is represented in yellow, while the human plasma protein is shown in cyan. Lipids and sugars are represented in lines and Ca^2+^ ions are shown in van der Waals. B) RMSD for the Ail/vitronectin complexed averaged over the 10 replicae. RMSF for the Ail (C) and Vitronectin (D) units in the protein/protein complex averaged over the 10 replicae. RMSD and RMSF for the individual replicae can be found in Supplementary Information (SI)

In this contribution we resort to all-atom molecular dynamics (MD) simulations performed on Ail embedded in a lipid bilayer mimicking the composition of the *Y. pestis* outer membrane, notably involving two dissymmetric leaflet the external one composed of *Y. pestis* lipopolysaccharides and the internal one composed of 74% PVPG, 21% PPPE, and 5% PVCL2.^44^ The initial structure of the bacterial protein has been retrieved from the pdb database (5JV8).^29^ Additional details on the set-up of the initial systems, which have been performed using the Charmm-gui web-server,^45^ may be found in Supplementary Information (SI). Human Vitronectin has also been modeled by unbiased MD simulation, starting from a structure retrieved by AlphaFold,^46^ note that coherently with the most probable oxidation state^33,38^ two sulfur bridges have been enforced between residues C156/C472 and C215/C293, respectively. Once stable structures of both proteins have been obtained protein/protein docking has been performed to identify the most probable interactions pathways between Vitronectin and the outer regions of Ail, using the online Haddock web server.^47^

On top of these initial structures, MD simulations of 800 ns each on ten independent replicas have been performed to assess the stability of the aggregate and the specific interactions taking place. Note that the structure of the unbound, transmembrane Ail, and of Vitronectin have also been modeled by 6 replicas. All the simulations have been performed using the NAMD code^48,49^ and analyzed using VMD.^50^ Proteins and lipids have been modeled using the charm force field,^51–53^ while water buffer is described by TIP3P.^54,55^ Additional details concerning the simulation strategy can be found in SI. The native Ail protein is composed of a narrow transmembrane β-barrel moiety, flanked by three more flexible arms: the N-terminal, the central, and the C-terminal one. The MD simulation of Ail embedded into a *Y. pestis* outer membrane shows a stable and rather rigid structure, as shown by the analysis of the time series of the root mean square deviation (RMSD) reported in Supplementary Information (SI). In contrast the globular and water-soluble vitronectin is composed by a core flanked by two disordered fragments, which in solution exhibit a higher flexibility.

As shown in Figure 1 A Ail and Vitronectin forms a persistent complex which locks the human protein at the surface of the bacterial outer membrane. As it can be appreciated from the Figure the two proteins exhibit an extended contact region. Interestingly, the latter appears embedded in the sugar moiety of the bacterial membrane. Not unexpectedly, the two proteins interact mostly through their flexible, or even disordered, regions, which involves notably the three arms of Ail. The high stability of the complex can also be appreciated by the evolution of the RMSD which plateaus rapidly at around 6 Å (Figure 1B) after equilibration and thermalization. Furthermore, the contribution of Ail to the total RMSD appears negligible compared to the one of Vitronectin. The more important flexibility of the human protein, as compared to its bacterial partner, can also be appreciated by the values of the per residues Root Mean Square Fluctuations (RMSF). Indeed, the more rigid Ail, and in particular its β-barrel core, leads to very small fluctuations comprised between 1.5 and 3.5 Å. Quite obviously, the peak of flexibility coincides with the arms and the extramembrane segments. Interestingly, and as it can be appreciated in SI the interaction with vitronectin only slightly perturbs the RMSF of the bacterial protein. This is also probably due to the fact that the flexible arms are still embedded in a dense and viscous environment, namely the outer bacterial membrane, which, thus, reduces its flexibility. Instead, while the core of Vitronectin is also rigids, a sharp peak can be observed for residues 100-150, which corresponds to the disordered region which is not in contact with Ail.

On the contrary, a very impressive flexibility reduction can be observed for the second disordered region (residues 350-450) which instead is in direct contact with the bacterial protein. Once again, this behavior is highly reminiscent of the one of intrinsically disordered proteins, which exploit their flexibility to act as hub in protein/protein interaction networks. As it will be also detailed further, the interaction with Vitronectin is not symmetric with respect to the tree arms of Ail, and indeed the N-terminal and the central arm appears to be much more involved in the stabilization than the C-terminal loop.

At a more residue-based resolution we may see from Table 1 that the protein/protein interaction is driven by a rather complex of both salt-bridges and Hydrogen bonds locking the two units together. However, their persistence along the whole MD simulation is also highly variable, and in some cases temporary interactions may be spotted, which globally contribute to the overall stabilization of the complex. In particular the interaction between R421 of Vitronectin and E55 of Ail appears to play a most particular role, being engaged in both a very persistent salt bridge (87% occurrence) and hydrogen bond (67%). This fact can also be inferred from the analysis of the distribution of the distances reported in Figure 2, which shows a very peaked, and highly gaussian residue-residue distance, indicative of a very stable and rigid arrangement. Furthermore, the interaction of D367 (Vitronectin) and D59 (Ail) is also highly persistent exceeding 75%, yet the hydrogen bond between the two residues appears less stable. This is also translated in a broader and less gaussian-shaped distribution of the residue-residue distance as shown in Figure 2.

**Figure 2.**
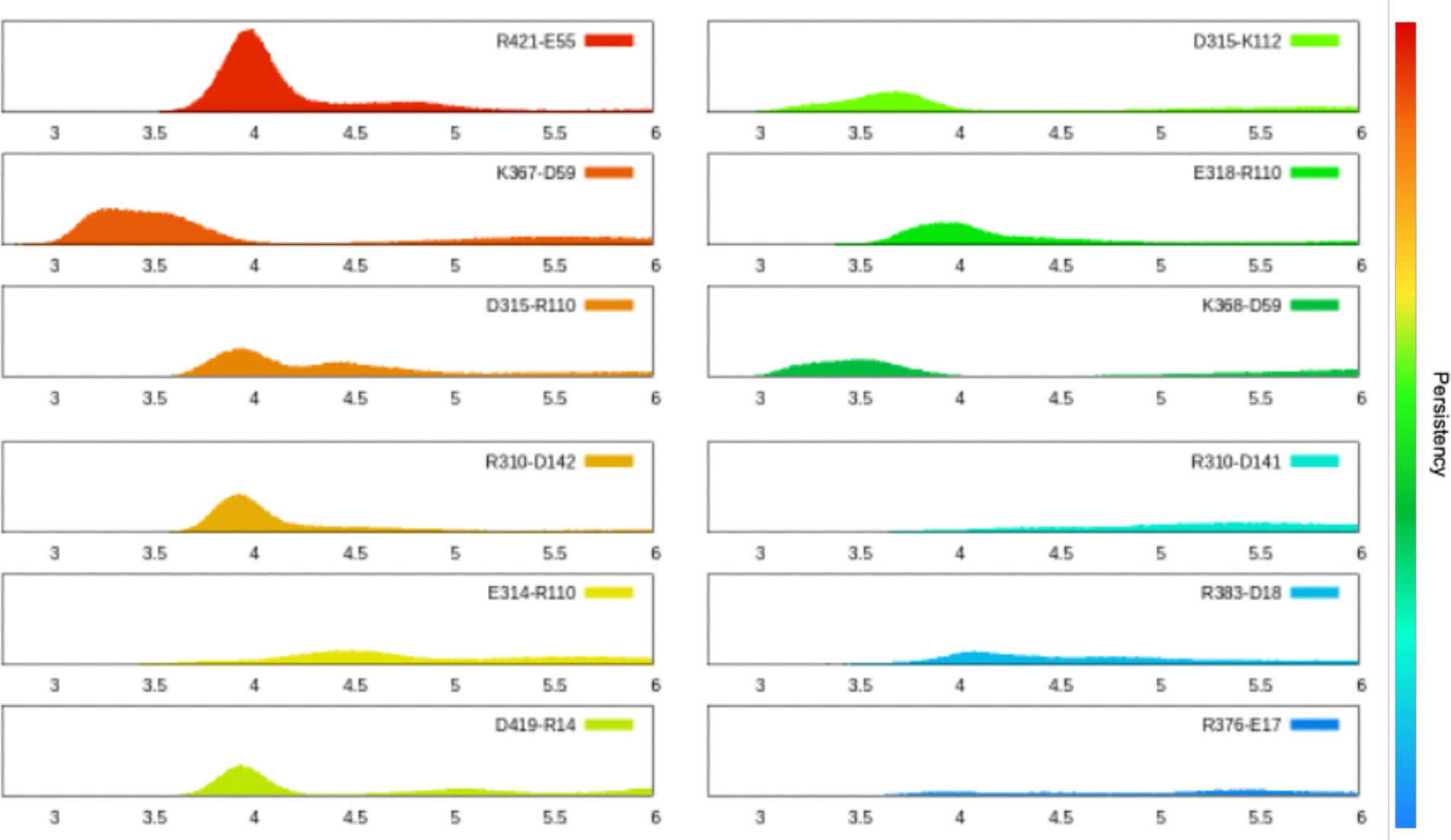
Distribution over the 10 replicae of the distance of the most persistent interactions highlighted in Table 1. Note that a color code identifying their importance in terms of persistence of the interaction is also used.

**Table 1.** Occurrences of the most persistent salt bridges and Hydrogen bonds takin g place at the Vitronectin/Ail interface averaged over the 10 replicae.

| **Vitronectin/Ail** | ***Salt Bridges*** | ***Hydrogen bonds*** |
| --- | --- | --- |
| *R421 – E55* | 87.6% | 66.64% |
| *K367 – D59* | 76.8% | 29.69% |
| *D315 – R110* | 53.9% | 34.25% |
| *R310 – D142* | 42.3% | 37.68% |
| *E314 – R110* | 40.8% | / |
| *D419 – R14* | 39.9% | 28.38% |
| *D315 – K112* | 39.3% | / |
| *E318 – R110* | 38.6% | 30.08% |
| *K368 – D59* | 38.6% | 23.12% |
| *R310 – D141* | 34.5% | / |
| *R383 – D18* | 33.2% | / |
| *R376 – E17* | 20.1% | / |
| *S312 – D142* | / | 35.18% |
| *R387 – D18* | / | 25.23% |

Other residues being strongly engaged in the protein/protein locking comprise D315 (Vitronectin) and R110 (Ail) having a persistency of about 54%, even if the distribution of the residue/residue distance presents some characteristic of a bimodal distribution. The latter bacterial residue is also interesting since it also leads to the interaction with E314 of Vitronectin. This interaction while having a persistency of about 40%, may lead to a synergic effect with the previous one, strongly reinforcing the protein/protein association. Finally, R310 (Vitronectin) and D18 (Ail) also lead to a highly persistent interaction which is also confirmed by the almost ideally gaussian distribution of the inter-residues distance. As shown in Figure 2, the interactions between the other residues listed in Table 1, leads to quite broad distributions indicative of a less important stabilization effect, with the only partial exception of D419 (Vitronectin)/R14 (Ail) which show a rather sharp peak at around 4 Å.

In Figure 3 the interprotein interactions whose persistence exceeds 40% are represented and mapped on the protein/protein interface region. As it was already surmised from the global structure, we may confirm that the bacterial protein participates differently in the stabilization of the complex. On the other hand, three main anchoring points can be evidenced, in which the N-terminal arms, in particular through the E55(Ail)/R421(Vitronectin), R14(Ail)/D419(Vitronectin), and D18(Ail)/R383(Vitronectin) network, clearly stands out as the main stabilizing factor of the complex.

**Figure 3.**
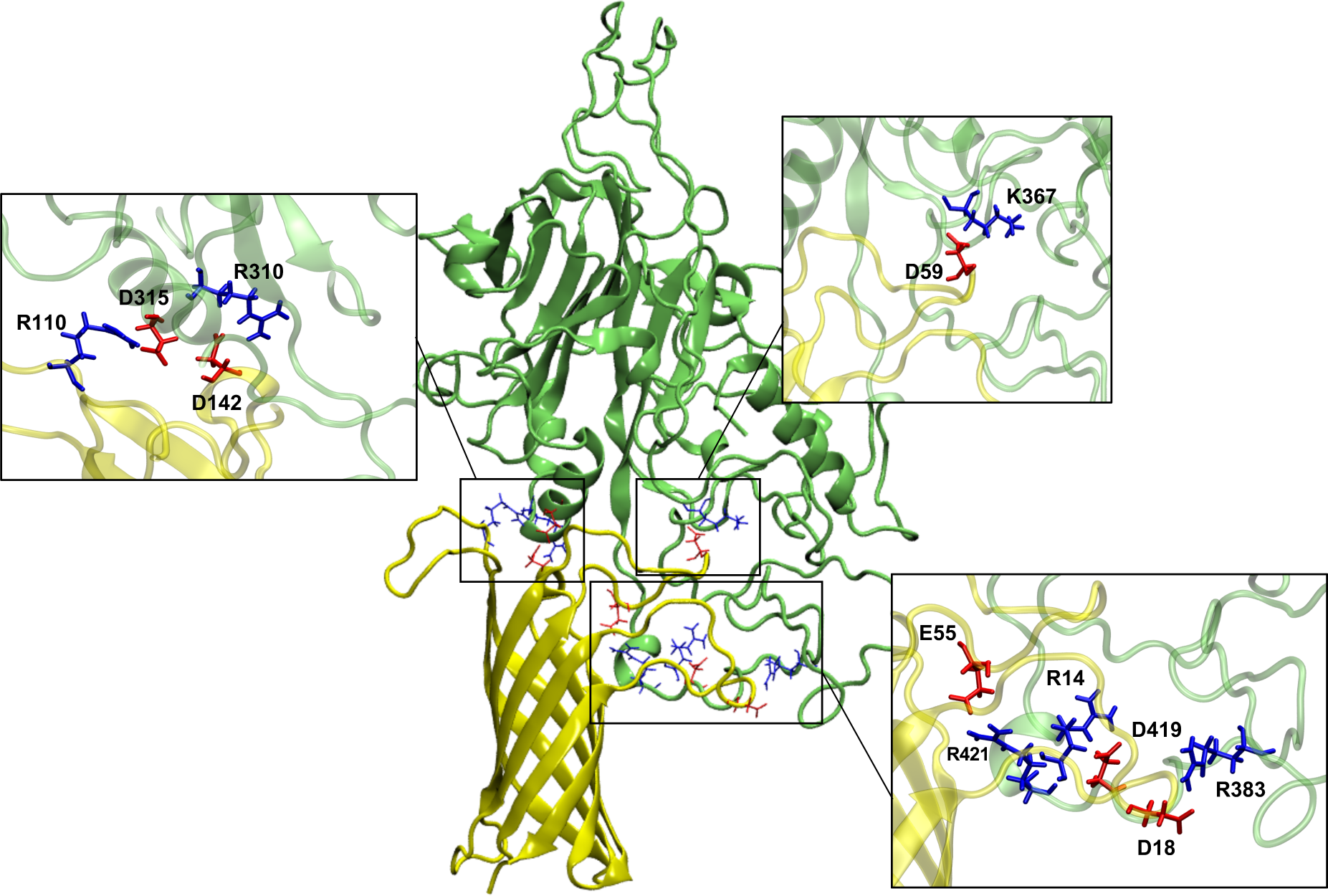
Location at the protein/protein interface of the most persistent interactions. In the inlays zooms showing the residues involved and their structural organizations are presented.

Due to the extended region of the protein/protein interaction it appears unlikely that a small molecule prodrug could efficiently destabilize the complex impeding Vitronectin recruitment. On the other hand, the possibility of targeting Ail and *Y. pestis* outer membrane with artificial peptides, specifically oriented towards the N-terminal loop to avoid the formation of the complex may appear as promising. In this respect, while the antibacterial peptide should be flexible and even disordered, its sequence should ideally contain both negative aminoacid, such as glutamate and aspartic acid, as well as arginine to favorably compete with the most important Vitronectin stabilizing interactions.

Finally, and as reported in Figure 4, we observed that one of the Ca^2+^ ions, which are largely present at the surface of the bacterial outer membrane, establishes persistent and very favorable interactions with solvent exposed Vitronectin residues. In particular, as shown in Figure 4B E221, E231, and Q216 appear as the most important factors leading to this specific interaction, which persists all along the sampling of our MD simulation.

**Figure 4.**
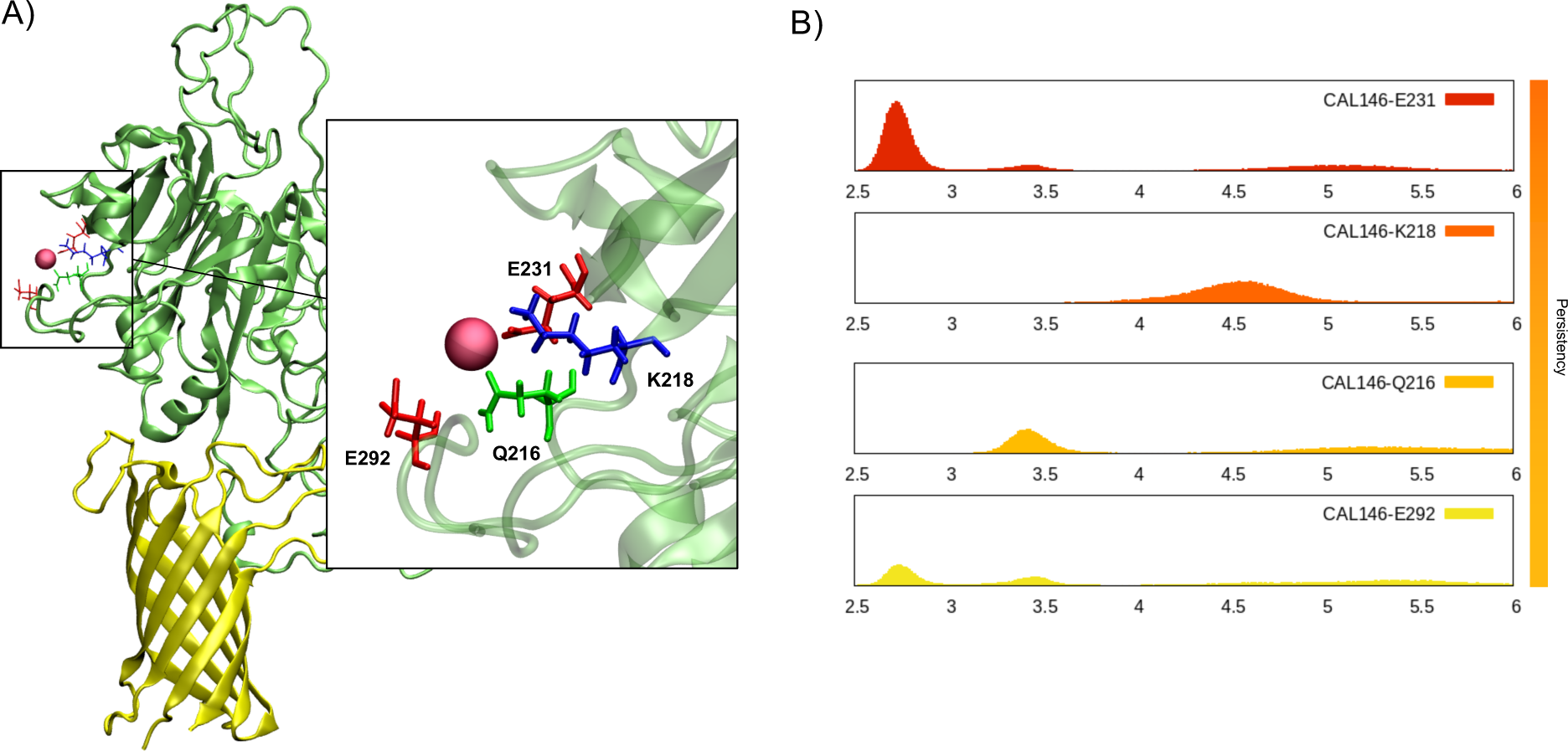
A) snapshot showing the persistent interaction of a Ca^2+^ ion with solvent exposed Vitronectin. The residues locking the ion are also represented in the inlay. B) Distribution of the distances between the ion and the closest vitronectin residues aver the whole 10 replicae.

Even if Ca^2+^ ions are present in the physiological medium, and specifically in plasma, their abundance is particularly important in the vicinity of the bacterial outer membrane. Therefore, this specific interaction may be regarded as an additional factor favoring the recruitment of Vitronectin at the bacterial interface.

Inhibiting the interaction between vitronectin and Ail paves the way to the development of novel antivirals, disrupting the protein complex and, thereby, restoring immune system functionality. It is noteworthy that the contacts between vitronectin and Ail are positioned on three main anchoring points, as presented in Figure 3. Indeed, since the seminal works of Easson and Stedman published in 1933,^56^ it is known that three interacting sites are necessary to assure biological recognitions.^57^

A drug design target, therefore, could be aimed at perturbing the protein-protein recognition by introducing new agents, generally low-molecular-weight organic molecules or peptides^58^ competing for the occupancy of the contact region. The goal of these agents is not to interact with all the three sites, as it is generally observed that inhibiting a single site is sufficiently effective in eliciting a biological response.^59^ In this regard, in a bacterial-targeted strategy, attention should be directed towards the identification of cavities on Ail, as it is established that small molecules, including peptides, generally insert into protein cavities.^59^ With the use of the FPocketWeb platform,^60^ we have determined that two of the three Ail/Vitronectin contact points correspond to druggable cavities as depicted in Figure S10. Namely they encompass the space between all the residues participating in the main electrostatic interactions already shown in Figure 3. Thus, these two cavities should clearly be prioritized as the most ideal target for a rational drug design endeavor.

In this letter we have presented the modeling of the interaction between the *Y. pestis* Ail protein, embedded in a model outer bacterial membrane, and the human Vitronectin. In particular our sampling largely exceeding the μs time-scale has confirmed the establishment of a persistent protein/protein complex, involving the interaction of the flexible regions of the human protein and the extramembrane loops of the bacterial partner. Interestingly, while the flexibility of Ail is only slightly affected by the binding, a much more pronounced effect can be observed for the human counterpart. Additionally, the individual residues which represent the most important driving forces for the formation of the complex have been identified. In particular, we have shown that the N-terminal and the central loop of the bacterial protein play a special role in the formation of the complex. Consequently, they also represent the most ideal druggable hot-spots. Yet, also considering the extension of the protein/protein interface, the use of antibacterial peptides, having disordered arrangement and negatively charged aminoacids, appears as the more promising to block Vitronectin interaction. The recruitment of Vitronectin at the bacterial surface being an important virulence factor, related to the evasion of the immune system, this strategy can be potentially interesting as an original strategy to counteract *Y. pestis* infection, especially in their most dangerous pulmonary and septicemic phases. The rational design of potential antibacterial peptides will be the object of a forthcoming contribution, yet our work enhances significantly the knowledge of the intricate coupling between human circulating proteins and bacteria in increasing the virulence of infections.

## ASSOCIATED CONTENT

### Supporting Information

Extended computational methodology. Evolution of the main interactions, hydrogen bonds and electrostatic, between Ail and Vitronectin and between Vitronectin a Ca^2+^ ion. Time evolution and distribution of the RMSD and RMSF for the individual replicae of all the studied systems. RMSD and RMSF of the flexible Ail arms in presence and absence of Vitronectin. Representation of the Ail druggable pockets. (file type, i.e., PDF).

## AUTHOR INFORMATION

### Corresponding Author

*F.B. A.M.

### Author Contributions

The manuscript was written through contributions of all authors. All authors have given approval to the final version of the manuscript.

## ACKNOWLEDGMENT

The authors thank GENCI and Explor computing centers and the Platform P3MB for computational resources. The authors thanks ANR and CGI for their financial support of this work through Labex SEAM ANR 11 LABEX 086, ANR 11 IDEX 05 02. The support of the IdEx “Université Paris 2019” ANR-18-IDEX-0001.

