## Supplementary Information for "Interaction Between *Yersinia pestis* Ail Outer Membrane Protein and the C-Terminal Domain of Human Vitronectin"

### COMPUTATIONAL METHODOLOGY

**Structure preparation.** All the initial models have been set up via the Charmm-Gui web server.[1–3] An initial system involving Ail embedded in a *Y. pestis* bacterial outer membrane model, has been constructed. The initial structure of Ail has been retrieved from PDB: 5JV8.[4] The composition of the bacterial outer membrane is highly asymmetric among the two leaflets. The internal one has been modeled considering randomly placing 200 lipids involving 74% PVPG, 21% PPPE, and 5% PVCL2. The external leaflet is composed of 200 *Y. pestis* lipopolysaccharides as defined by the Charmm-Gui default parameters.[2] Note that a water buffer has been used for each membrane systems and in all cases a physiological salt (KCl) concentration of 0.15 M has been enforced. The outer leaflet of the *Y. pestis* outer membrane being composed of lipopolysaccharides whose negative charges are neutralized by $Ca^{2+}$ ions, these cations have been added in vicinity of the outer polar head region by Charmm-Gui. The solvated C-terminal domain of Vitronectin has been modeled starting from the AlphaFold model,[5,6] which has been solvated in a water box bearing a physiological (0.15 M) concentration of KCl using again the Charmm-gui web server.

Charmm36m force field and its extension for lipids and bacterial lipids[7–9] have been consistently used for all the systems.

After a first equilibration of the two systems representative snapshots of membrane embedded Ail and solvated Vitronectin have been extracted. The two proteins have been docked using the Haddock web-server[10] to obtain suitable initial conformations of the protein/protein complex. The former has then been embedded in a *Y. pestis* outer membrane having the same composition as for the Ail system using again the Charmm-Gui web interface with the same protocol as detailed before.

**Computational details.** All the calculations were performed using NAMD.[11,12] Every simulation was performed using hydrogen mass repartition (HMR),[13] allowing to use a time step of 4 fs for each calculation. All the simulations have been performed in the isothermal and isobaric (NPT) ensemble using the Langevin thermostat[14] and barostat[15] to enforce conservation of the temperature and pressure, respectively, respectively. The initial system was first minimized by 50000 steps and subsequently equilibrated and thermalized by progressively removing constraints on the protein backbone atoms during 36 ns. Lipids have not been constrained to avoid the formation of artificial water pores at the box borders. As previously stated, three different systems have been built, namely the Ail/Vitronectin complex embedded in the bacterial membrane, Ail embedded in the bacterial membrane, and solvated Vitronectin.

The system involving the Ail/Vitronectin complex has modeled considering 10 independent replicae, and each one has been propagated for about 880 ns, i.e. with an average of 34 million steps for each trajectory, leading to a total simulation time of about 9 μs. In the case of Ail embedded in the lipid membrane, six independent replicae have been considered, and each one has been propagated for about 460 ns with an average of 22 million steps for each trajectory. Finally, solvated Vitronectin, has also been modeled considering 6 independent replicae, propagated for about 880 ns with an average of 22 million steps of trajectory each. As a matter of fact, we choose to have longer simulation times for vitronectin due to the necessity to correctly sampling the extended disordered loops flanking the rigid protein core.

**Analysis.** Cpptraj[16] was employed for processing trajectory output files, performing Root Mean Square Deviation (RMSD) and Root Mean Square Fluctuation (RMSF) analyses. Gnuplot software was utilized to generate the time series and distribution plots.

The trajectories have been visualized and analyzed using VMD,[17] which was also employed to identify hydrogen bonds throughout the trajectory, applying a distance cutoff of 3 Å and an angle cutoff of 20°. For the recognition of electrostatic interactions, charged atoms within a distance of 6 Å from each other were identified using VMD. This specifically involved arginine, lysine, as well as glutamic and aspartic acid residues. Subsequently, the percentage of the presence of these interactions along the whole simulation was calculated using Cpptraj in addition to a custom Python code.

| | | Rep1 | Rep2 | Rep3 | Rep4 | Rep5 | Rep6 | Rep7 | Rep8 | Rep9 | Rep10 | Mean |
|---|---|---|---|---|---|---|---|---|---|---|---|---|
| **R421** | **E55** | 59,47% | 40,76% | 84,78% | 98,61% | 102,55% | 73,83% | 52,34% | 72,11% | 73,85% | 8,14% | **66,64%** |
| **R310** | **D142** | 28,16% | 10,02% | 80,56% | 56,44% | 36,47% | 1,02% | 73,36% | 74,19% | 5,76% | 10,83% | **37,68%** |
| **S312** | **D142** | 29,81% | 21,45% | 61,39% | 29,54% | 3,57% | 42,98% | 72,44% | 62,65% | 14,03% | 13,97% | **35,18%** |
| **R110** | **D315** | 12,67% | 83,49% | 29,21% | 67,04% | 6,95% | 3,80% | 46,64% | 41,18% | 14,26% | 37,22% | **34,25%** |
| **R110** | **E318** | 89,51% | 7,50% | 21,05% | 4,68% | 0,23% | 87,82% | 70,77% | 19,07% | 0% | 0,18% | **30,08%** |
| **K367** | **D59** | 9,42% | 36,28% | 42,00% | 26,16% | 29,89% | 46,55% | 36,29% | 9,97% | 40,04% | 20,25% | **29,69%** |
| **R14** | **D419** | 8,50% | 11,16% | 7,61% | 107,04% | 8,06% | 0% | 2,13% | 105,40% | 0% | 33,80% | **28,38%** |
| **R387** | **D18** | 6,46% | 74,38% | 65,52% | 36,71% | 11,21% | 7,97% | 0% | 35,13% | 14,91% | 0% | **25,23%** |
| **K368** | **D59** | 43,01% | 4,76% | 54,33% | 25,83% | 37,21% | 8,71% | 15,41% | 24,56% | 4,74% | 12,65% | **23,12%** |

**Table S1.** Percentages of presence of hydrogen bonds between AIL and Vitronectin for each one of the ten replicas as well as their average. A color code highlighting the persistence of the different interactions is used: contacts exceeding 60% are represented in red, those surpassing 40% in orange, those comprised between 20% and 40% in yellow. Interactions having a persistence time lower than 20% are represented in blue, and under 7% in blue. The aminoacids identified on the first column belong to vitronectin and on the second column to Ail.

| | | Rep1 | Rep2 | Rep3 | Rep4 | Rep5 | Rep6 | Rep7 | Rep8 | Rep9 | Rep10 | Mean |
|---|---|---|---|---|---|---|---|---|---|---|---|---|
| **R421** | **E 55** | 100,0% | 80,4% | 100,0% | 99,3% | 98,0% | 95,3% | 93,2% | 99,3% | 99,7% | 11,2% | **87,6%** |
| **K367** | **D59** | 43,9% | 91,9% | 99,4% | 85,4% | 64,4% | 95,4% | 82,2% | 61,4% | 88,8% | 54,8% | **76,8%** |
| **D315** | **R110** | 67,3% | 92,9% | 50,1% | 88,5% | 15,5% | 16,1% | 86,2% | 66,9% | 13,4% | 42,2% | **53,9%** |
| **R310** | **D142** | 26,2% | 11,8% | 78,6% | 56,3% | 65,3% | 0,9% | 74,0% | 73,8% | 13,4% | 22,3% | **42,3%** |
| **E314** | **R110** | 53,5% | 34,4% | 74,2% | 34,2% | 5,5% | 17,6% | 74,9% | 74,8% | 14,0% | 25,0% | **40,8%** |
| **D419** | **R14** | 79,7% | 12,7% | 26,1% | 96,1% | 14,8% | 5,5% | 22,8% | 86,2% | 18,5% | 36,3% | **39,9%** |
| **D315** | **K112** | 56,4% | 13,6% | 69,4% | 6,0% | 0,5% | 96,6% | 83,2% | 67,0% | 0,6% | 0,1% | **39,3%** |
| **E318** | **R110** | 94,9% | 12,2% | 52,7% | 10,3% | 0,7% | 96,5% | 80,5% | 38,1% | 0,2% | 0,4% | **38,6%** |
| **K368** | **D59** | 93,3% | 16,8% | 15,1% | 36,4% | 76,1% | 51,5% | 38,9% | 12,9% | 14,4% | 30,3% | **38,6%** |
| **R310** | **D141** | 21,4% | 15,3% | 59,1% | 50,7% | 4,4% | 18,5% | 63,5% | 65,8% | 17,6% | 28,8% | **34,5%** |
| **R383** | **D18** | 53,6% | 0,0% | 59,4% | 2,9% | 5,5% | 68,4% | 94,7% | 23,5% | 2,0% | 22,0% | **33,2%** |
| **R376** | **E17** | 34,5% | 83,5% | 28,6% | 0,2% | 4,3% | 3,2% | 7,8% | 37,2% | 1,5% | 0,5% | **20,1%** |

**Table S2.** Percentages of presence of electrostatic interactions between AIL and Vitronectin for each one of the ten replicas as well as their average. A color code highlighting the persistence of the different interactions is used: contacts exceeding 60% are represented in red, those surpassing 40% in orange, those comprised between 20% and 40% in yellow. Interactions having a persistence time lower than 20% are represented in blue. and under 7% in blue. The aminoacids identified on the first column belong to vitronectin and on the second column to Ail.

| | | Rep1 | Rep2 | Rep3 | Rep4 | Rep5 | Rep6 | Rep7 | Rep8 | Rep9 | Rep10 | Mean |
|---|---|---|---|---|---|---|---|---|---|---|---|---|
| **CAL 146** | **E 231** | 27,07% | 56.37% | 28.85% | 56.96% | 54.68% | 93.01% | 13,00% | 85.60% | 27.79% | 16.14% | **45,98%** |
| **CAL 146** | **K 218** | 23.27% | 51.91% | 26.17% | 51.77% | 51.60% | 6.96% | 10.61% | 72.46% | 22.50% | 15.40% | **33,27%** |
| **CAL 146** | **Q 216** | 25.03% | 45.16% | 24.75% | 41.73% | 52.52% | 24.69% | 8.87% | 62.48% | 19.36% | 14.11% | **31.87%** |
| **CAL 146** | **E 292** | 13.28% | 13.87% | 14.26% | 34.04% | 15.49% | 96.35% | 0.48% | 57.01% | 8.93% | 6.89% | **26,05%** |

**Table S3.** Percentages of presence of distances below 6Å between vitronectin aminoacids and the closest $Ca^{2+}$ ion, i.e. residue 146, for each ten replicas and their average. A color code highlighting the persistence of the different interactions is used: contacts exceeding 60% are represented in red, those surpassing 40% in orange, those comprised between 20% and 40% in yellow. Interactions having a persistence time lower than 20% are represented in blue. and under 7% in blue.

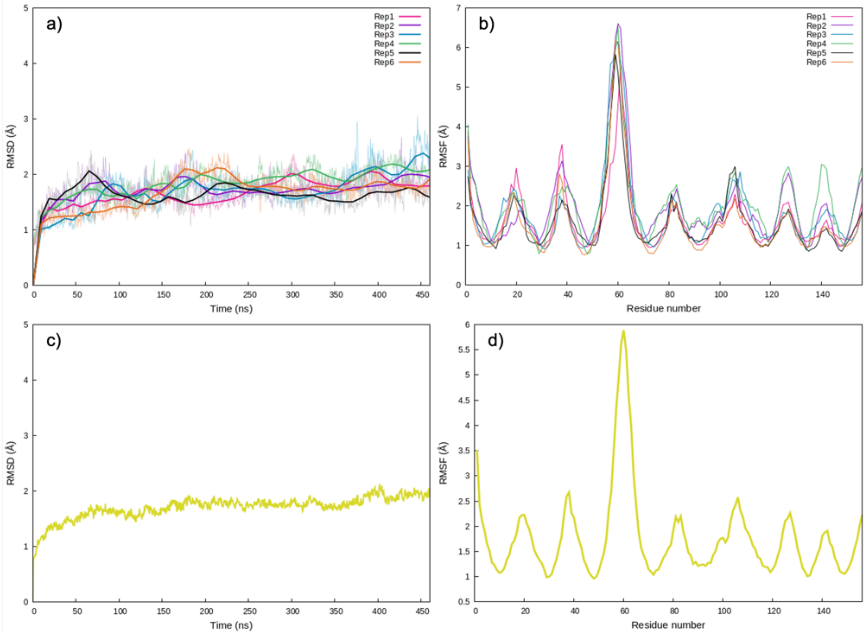


**Figure S1.** Time evolution of the RMSD of Ail for each of the 6 replicae a) and the corresponding RMSF b). Average of the RMSD over the 6 replicae c) and of the RMSF d).

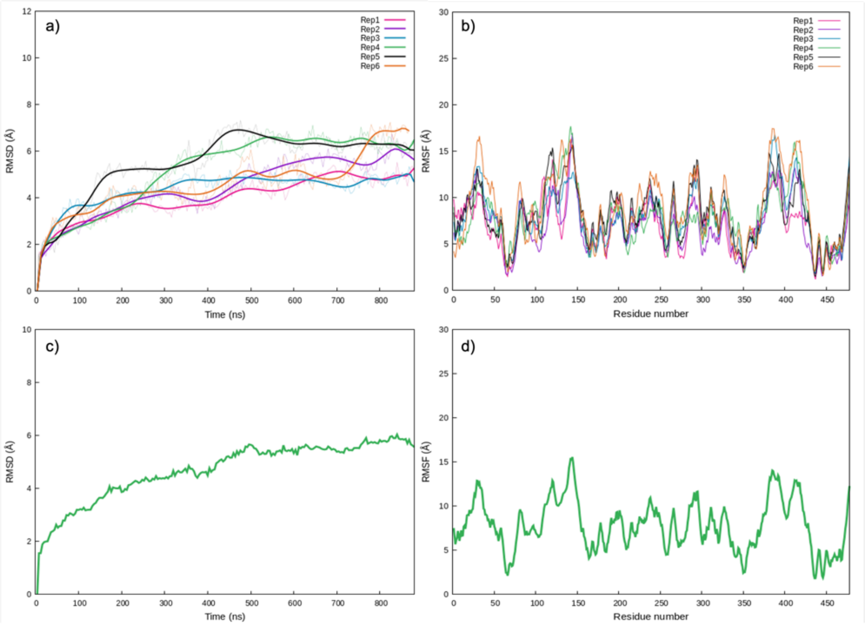


**Figure S2**. Time evolution of the RMSD of solvated Vitronectin for each of the 6 replicae a) and the corresponding RMSF b). Average of the RMSD over the 6 replicae c) and of the RMSF plicae.

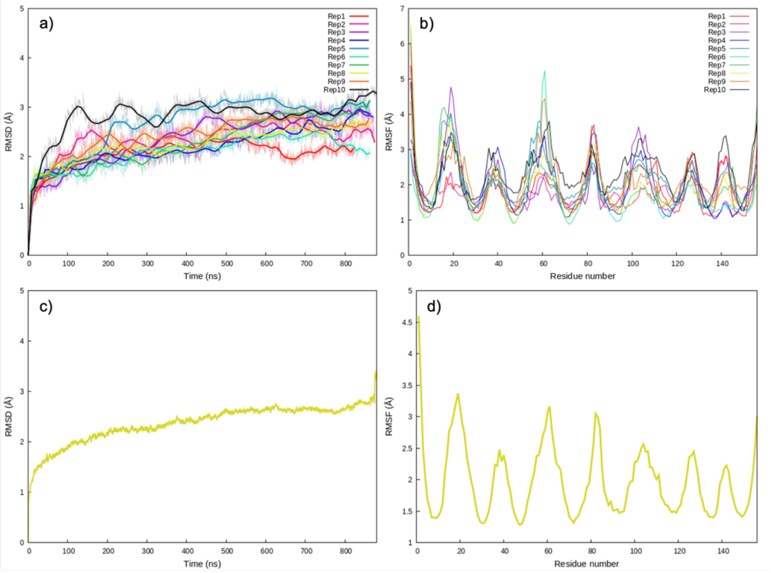


**Figure S3.** Time evolution of the RMSD of the Ail fragment in the Ail/Vitronectin complex of each of the 10 replicae a) and the corresponding RMSF b). Average of the RMSD over the 10 replicae c) and of the RMSF d).

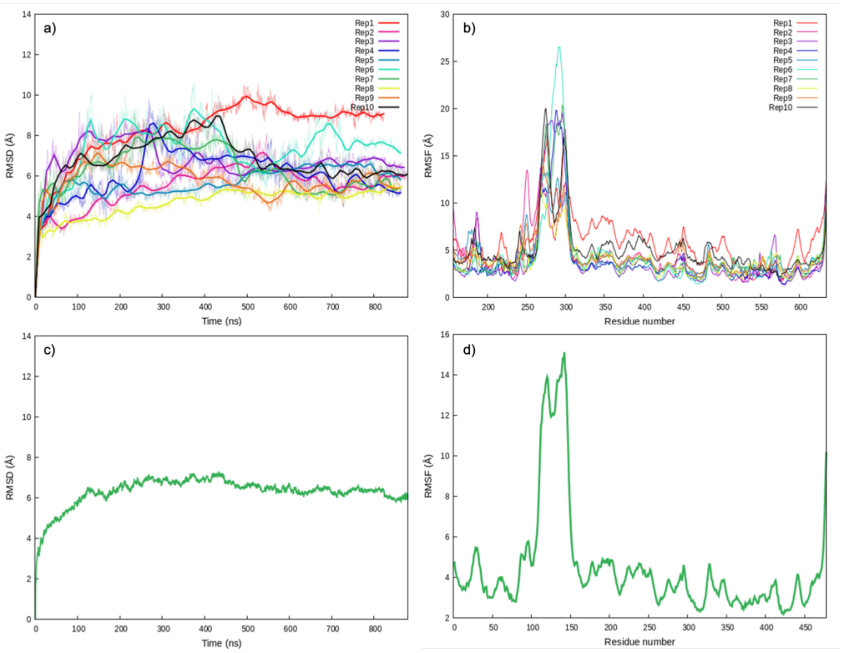


**Figure S4.** Time evolution of the RMSD of the Vitronectin fragment in the Ail/Vitronectin complex of each of the 10 replicae a) and the corresponding RMSF b). Average of the RMSD over the 10 replicae c) and of the RMSF d).

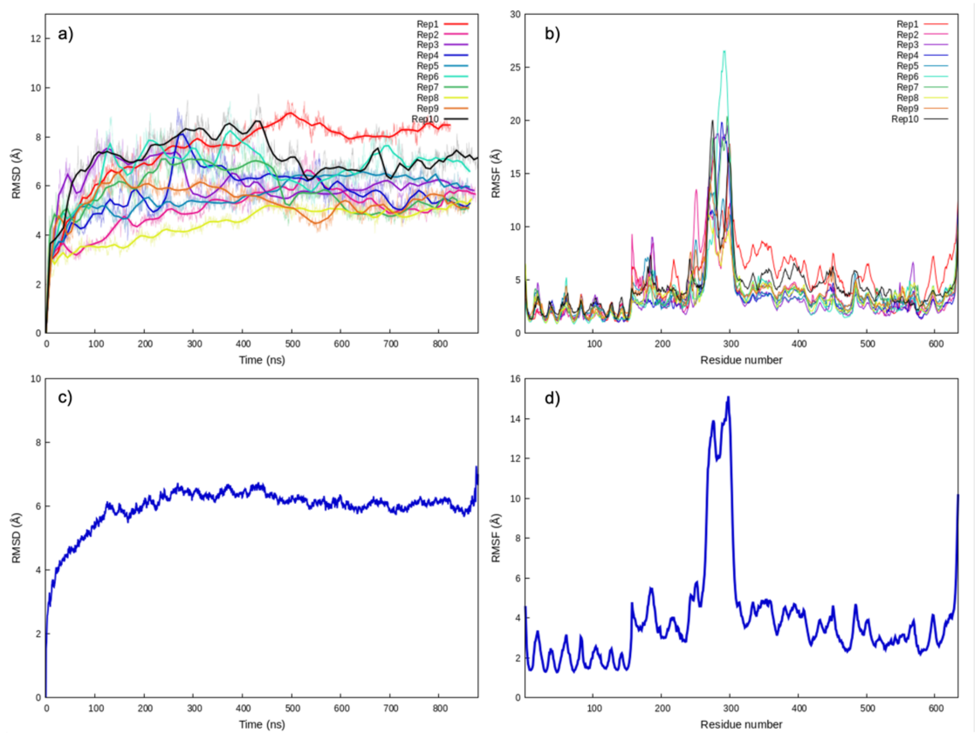


**Figure S5.** Time evolution of the RMSD of the total Ail/Vitronectin complex of each of the 10 replicae a) and the corresponding RMSF b). Average of the RMSD over the 10 replicae c) and of the RMSF d).

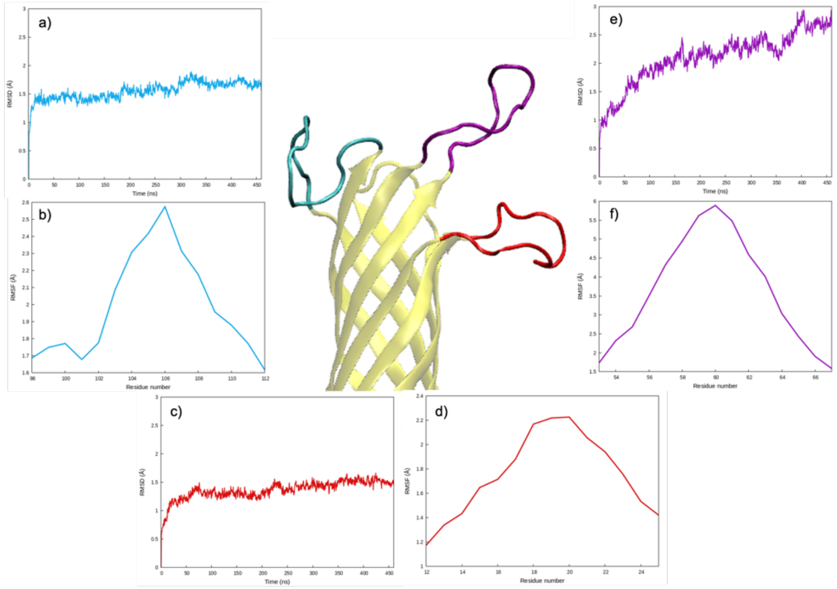

**Figure S6.** Flexibility of the Ail Arms for the membrane embedded Ail averaged over the 6 replicae. a) RMSD of ‘arm 3’ (residue 98 to 112) and b) RMSF, c) RMSD of ‘arm 1’ (residue 12 to 25) and d) RMSF, e) RMSD of ‘arm 2’ (residue 53 to 67), and f) RMSF. The arms are also evidenced in the reproduced snapshots using the same color code as for the curves, i.e. blue for ‘arm 3’, red for ‘arm 2’, and purple for ‘arm 1’.

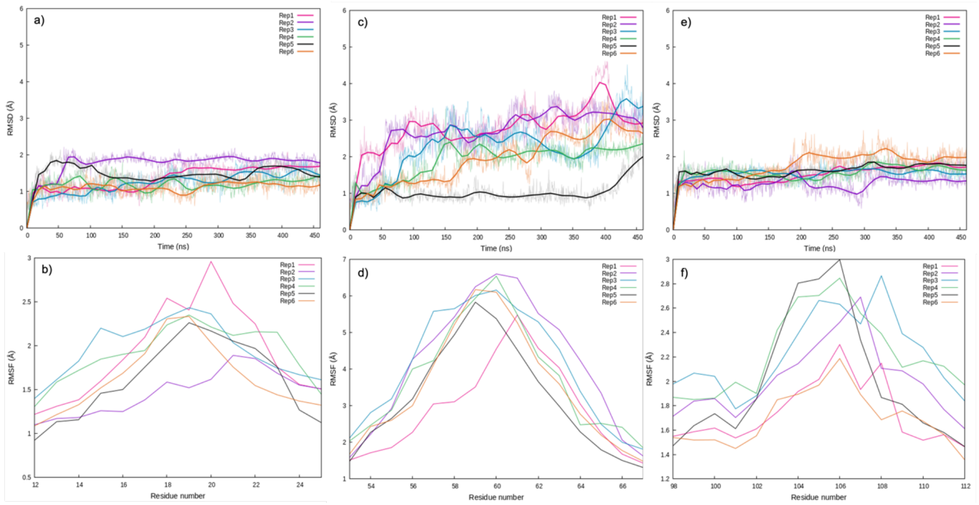

**Figure S7.** Flexibility of the Ail arms in the membrane embedded system represented for each replica. RMSD of ‘arm 1’ (residue 12 to 25), b) RMSF of ‘arm 1’, c) RMSD of ‘arm 2’ (residue 53 to 67), d) RMSF of ‘arm2’, e) RMSD of ‘arm 3’ (residue 98 to 112). f) RMSF of ‘arm 3’.

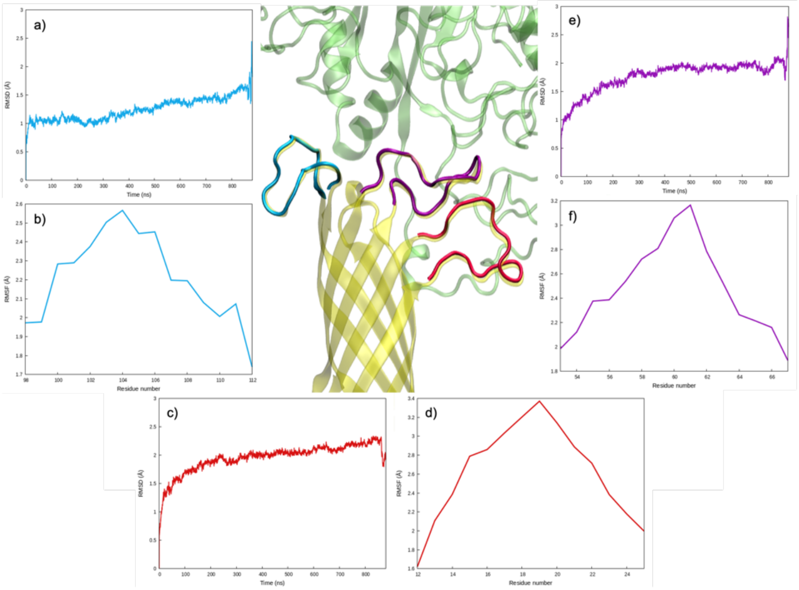


**Figure S8.** Flexibility of the Ail Arms for the Ail/Vitronectin complex averaged over the 10 replicae. a) RMSD of 'arm 3' (residue 98 to 112) and b) RMSF, c) RMSD of 'arm 1' (residue 12 to 25) and d) RMSF, e) RMSD of 'arm 2' (residue 53 to 67), and f) RMSF. The arms are also evidenced in the reproduced snapshots using the same color code as for the curves, i.e. blue for 'arm 3', red for 'arm 2', and purple for 'arm 1'.

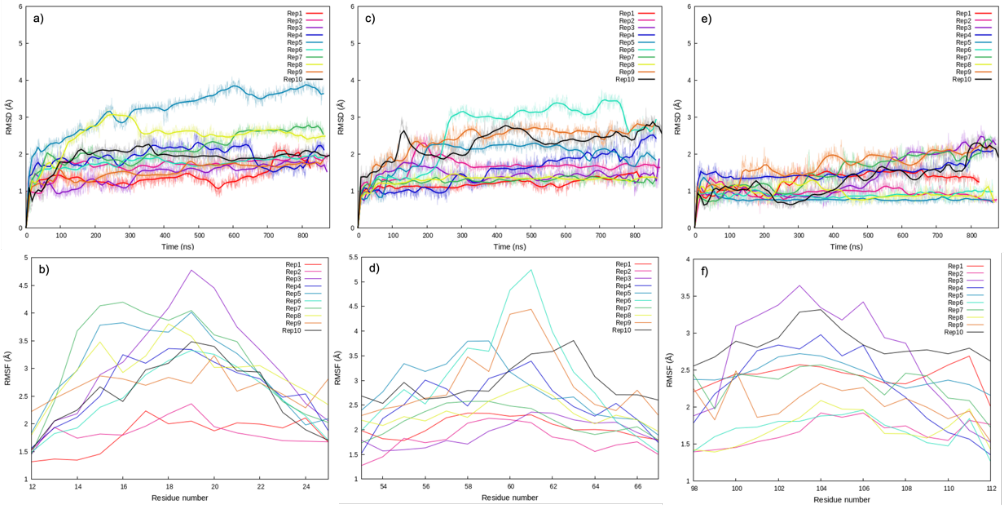


**Figure S9.** Flexibility of the Ail arms in the membrane embedded system represented for each replica. RMSD of 'arm 1' (residue 12 to 25), b) RMSF of 'arm 1', c) RMSD of 'arm 2' (residue 53 to 67), d) RMSF of 'arm2', e) RMSD of 'arm 3' (residue 98 to 112). f) RMSF of 'arm 3'.

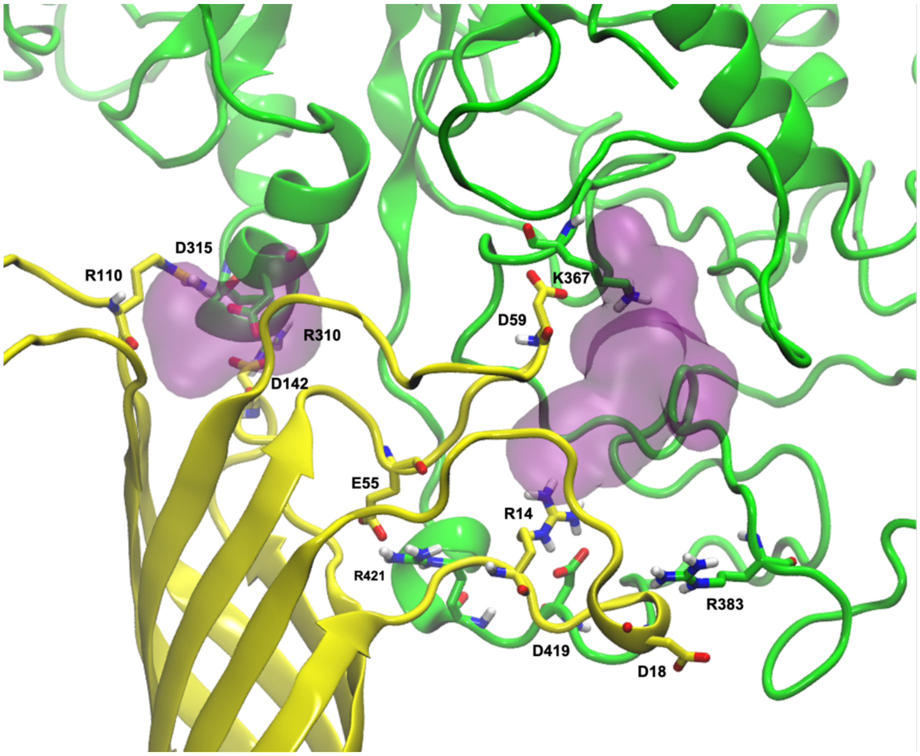


**Figure S10.** Visualization of druggable cavities in the Ail/Vitronectin complex.
